# Unpacking Chromatin Accessibility with Fiber-seq

**DOI:** 10.64898/2026.08.14.744917

**Authors:** Kerry L. Bubb, Molly Perchlik, Joshua T. Cuperus, Christine Queitsch

## Abstract

Chromatin accessibility has long been used as a marker for regions of DNA with regulatory potential. Fiber-seq detects chromatin accessibility on individual DNA fibers, enabling analyses beyond the identification of the accessible chromatin regions (ACRs). By providing single molecule level high resolution, Fiber-seq provides unprecedented qualitative descriptions, including potential categorizations of ACRs, identification of internal transcription factor footprints and nucleosome positioning within individual DNA fibers. As with all tools, the power of this technique depends on careful experimental design and data analysis -- incorrect usage will result in incorrect conclusions. Here we offer guidelines and flag potential pitfalls when generating and analyzing Fiber-seq data, such as (1) the optimum levels of adenosine methylation per-fiber, (2) the power of per-fiber state inference, (3) the importance of controlling for read depth and methylation rates when comparing across samples, (4) the limitations of long-read sequence mapping, and (5) suggestions for identification of differentially accessible peaks across samples.

## Introduction

Enzymes like DNase I, Tn5, and Hia5, which preferentially modify non-nucleosome-bound DNA, have proven invaluable to delineate genomic regions accessible to regulatory elements and infer protein-DNA interactions that govern regulation of gene expression. For the first few decades, chromatin accessibility assays were performed on bulk tissue, with accessibility defined as a binary state. However, even with these older accessibility assays, it was apparent that there were differences in both degree and quality of accessible states. The development of methods to purify single cell-types (Deal & Henikoff 2011) and subsequent advancements in single-cell profiling technologies (Baek & Lee 2020; Wang et al. 2025) indicated that this variability was not simply due to a mixture of cell types, but an inherent property of chromatin accessibility.

Fiber-seq (Stergachis et al. 2020) outperforms prior methods by measuring chromatin accessibility on long (>15 kb) single molecules at much higher per-bp resolution than previous methods. Compared with ATAC-seq, Fiber-seq offers three major advantages. First, it enables the analysis of genomic regions that cannot be mapped using short reads, such as long repetitive regions. Second, it reveals the existence of multiple classes of ACRs across the genome, including those with short, irregularly positioned regions of accessibility on underlying single molecules and others with longer, consistently positioned ones. Third, it reveals high-resolution details about individual ACRs, for example enabling the detection of transcription factors and nucleosome footprints on each underlying molecule. Recent studies suggest that ACRs that appear unchanging at an accessibility level can nonetheless change dramatically in terms of transcription factor occupancy (Hu et al. 2025). However, this higher resolution also increases the potential for new artifacts. For example, within a single Fiber-seq experiment, the degree of adenosine methylation varies widely across fibers. In addition, the distribution of per-fiber methylation often differs between Fiber-seq experiments. These technical differences must be accounted for to avoid conflating them with biologically meaningful differences.

Previously, we characterized the accessible chromatin landscape of the maize reference inbred strain B73 (Bubb et al. 2025) using the Fiber-seq method (Stergachis et al. 2020) and associated tools. We described a variety of types of accessible regions, discovered hundreds of individual LTR retrotransposons with persistent internal promoter and enhancer accessibility, and showed that regions with diffuse chromatin accessibility were more likely to be infiltrated by transposons. We are now in the process of extending our analysis to compare accessibility across trios consisting of two inbred parental lines and their F1 offspring. Our aim is to determine how chromatin accessibility patterns associated with a given genome in a particular tissue change in a different trans-environment. By comparing the chromatin accessibility patterns of the parental inbreds with the F1 hybrid, we also aim to gain insights into the mechanism of hybrid vigor. This task presents new challenges. Unlike with human samples, in which the two genomes (haplotypes) in each diploid sample are methylated in the same experiment and then computationally resolved (Vollger et al. 2025), we can prepare essentially single genomes in inbred parental lines and compare them to that same entire genome in the F1 to understand the influence of trans-acting factors on chromatin accessibility.

In this paper, we will (i) highlight how sample preparation quality interacts with important properties of the analytical algorithms and can influence the conclusions drawn, (ii) point out potential sources of false negatives when attempting to identify regions of differential accessibility across samples, and (iii) propose strategies to identify regions with truly differential accessibility across samples. We will not be describing the fibertools and FIRE algorithms that enable initial processing of raw Fiber-seq data, as those details are published elsewhere (Jha et al. 2024; Vollger et al. n.d.). In many ways, this manuscript is similar to our previous analysis of the properties of ATAC-seq (Bubb & Deal 2020). By outlining common pitfalls and suggesting practical recommendations, the paper aims to promote Fiber-seq as a powerful tool to advance our understanding of the dynamics of chromatin accessibility across diverse biological systems.

### 1. Initial metrics for evaluating Fiber-seq sample quality differ from those used in short-read accessibility assays

Like ATAC-seq, Fiber-seq is used to identify genomic positions of accessible chromatin regions (ACRs). Unlike ATAC-seq, however, Fiber-seq also generates rich accessibility descriptions of the long individual DNA molecules contained within a single experiment (Vollger et al. n.d.). A user new to Fiber-seq might first be tempted to examine whether it is feasible to simply bulk the raw signal (m6A in this case) and call peaks as is done with previous end-sequencing methods, such as MNase-seq, DNAse-seq, ATAC-seq. We therefore wish to highlight why this is not a good approach and why the fiber-level state inferences are essential for proper peak calling. Furthermore, because of the differences between the two assays, it is not appropriate to use the same quality metrics when evaluating newly prepared samples.

End-sequencing chromatin accessibility methods rely on the generation and sequencing of small fragments (generally ∼200 bps) where each end of the fragment is accessible to the enzyme used in the assay (**Fig 1A**, top). The enzyme used in Fiber-seq creates single nucleotide modifications which are then detected via long-read sequencing, and thus detection of one modification does not depend on the presence of another one nearby (**Fig 1B**, top). In end-based assays the first raw data that we can directly observe is the per-base count of ends, most of which are concentrated in regions where two ends are approximately 200 bp apart (**Fig 1A**, middle). In Fiber-seq, the first raw data that we can directly observe is methylated adenosines on individual DNA fibers, some of which are in >200bp accessible regions and some of which are in linker spaces between nucleosomes (**Fig 1B**, middle). If we simply make a histogram of the raw data from each assay within this 5-kb window, the end-sequencing data produces much cleaner peaks of accessibility than Fiber-seq (**Fig 1A&B**, bottoms).

**Figure 1.**
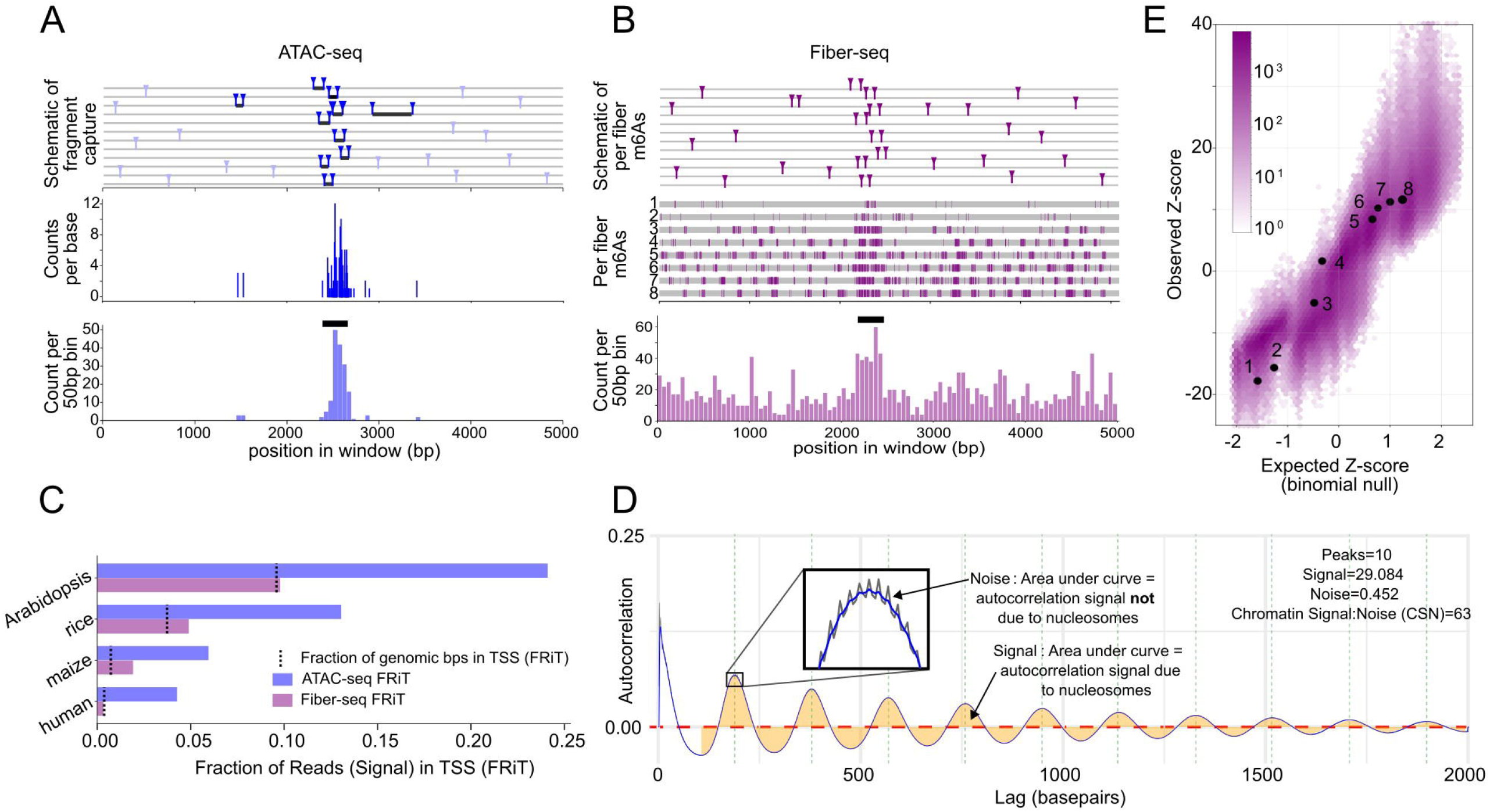
Properties of raw data obtained in Fiber-seq compared to that obtained with end-seq methods. **(A)** Workflow of end-seq based accessibility assays, such as MNase-seq, DNase-seq, and ATAC-seq. This example is taken from an ATAC-seq sample [B73_chr08:76,262,000-76,267,000]. Schematic of enzymes cleaving DNA at accessible sites (top). Pairs of cleavage sites resulting in PCR amplified and end-sequenced fragments are in dark blue; singleton cleavage sites not resulting in end-sequencing are shown in light blue. Raw observed data (middle). Histogram of raw observed data using non-overlapping 50 bp bins, with the black rectangle indicating an Accessible Chromatin Region (bottom). **(B)** Workflow of the Fiber-seq accessibility assay [LH287_ptg000001l:1,145,000-1,150,000]. Schematic of enzyme methylating adenosines at accessible sites (top). Raw observed data (middle). A histogram of raw observed data using non-overlapping 50-bp bins, with the black rectangle indicating an Accessible Chromatin Region (bottom). **(C)** FRiT score for pairs of ATAC- and Fiber-seq samples from different species, with FRiT defined as the amount of signal occurring in the 400 bps upstream of a Transcription Start Site divided by the total amount of signal. Arabidopsis (Col-0): ATAC-seq and Fiber-seq sample is in-house; rice (Kitaake): ATAC-seq is from (Swift et al. 2023), Fiber-seq sample is Bing (OSat/Kt-HT); maize (B73): ATAC-seq is from (Marand et al. 2025), Fiber-seq sample is from (Bubb et al. 2025); human (GM12878): ATAC-seq samples is from Groza et al. (2023), Fiber-seq data is from Vollger et al. (n.d.). **(D)** An autocorrelation plot of m6As within 2 kb. Peaks indicate inter-nucleosome linker positions; troughs indicate nucleosome locations. The area under the smoothed curve (indicated in gold) provides the autocorrelation signal due to nucleosomes. The area between the actual curve in dark gray and the smoothed curve in blue indicates the autocorrelation signal not due to nucleosomes (noise). Vertical green dotted lines indicate local maxima in the smoothed curve. **(E)** Superimposed Q-Q plots of the normalized m6A counts on fibers in 50,000 5-kb windows. Fibers in the 5-kb window shown in (B, middle) are in red, with numbered fibers labeled.

Because Fiber-seq generates single-nucleotide signal (m6A) in inter-nucleosome linkers, metrics that are typically used to report sample quality, such as a Fraction of Reads in Peaks (FRiP) or the less peak-caller-sensitive Fraction of Reads in TSSs (FRiT), do not accurately reflect the utility of the sample (**Fig 1C**). For example, the human sample shown (GM12878) was used to develop the FIRE pipeline (Vollger et al. n.d.) and yet, in aggregate, has no more signal in the TSS region than expected due to chance. Although some samples do have FRiP scores substantially exceeding that expected due to chance, notably the previously published maize sample (Bubb et al. 2025), optimizing only for FRiP score favors samples with very low levels of methylation and would result in an inability to detect other important features, such as nucleosomes and other protein footprints.

Another metric that can be derived from raw data alone is whether the spacing between methylated adenosines on the same fiber indicates the presence of nucleosomes. This metric has the added advantage that it requires very few reads -- typically just a few thousand -- and so can be performed with small sample quantities prior to expensive deep-coverage sequencing. We use an autocorrelation plot of all methylated adenosines within a distance (Lag) of 2 kb to visualize nucleosome and linker positions, the local minima and maxima, respectively (**Fig 1D**). The amount of vertical fluctuation in the autocorrelation plot due to nucleosome positioning can be defined as the nucleosome ‘Signal’ (area between the smoothed autocorrelation curve and the y=0 line), and the vertical fluctuation due to anything else is the ‘Noise’ (area between the observed and smoothed autocorrelation curves, seen in inset, **Fig 1D**). The ratio between these two is called the Chromatin Signal-to-Noise (CSN). We have found that a CSN ratio above 25 indicates that the sample contains both intact chromatin and sufficient adenosine methylation. Because identifying nucleosomes is secondary to our primary goal of identifying accessible regions, the aim should not be to maximize this metric but simply be above the aforementioned threshold.

Finally, a feature of the data that can be seen with Fiber-seq but not with bulk end-sequencing methods is that, within a sample preparation, the per-fiber adenosine methylation rate deviates greatly from a null expectation that the Hia5 methyltransferase is methylating adenosines on each fiber with equal probability -- even at the same genomic location. Although there is no way of knowing the range of transposition across different genomes/cells in bulk ATAC-seq experiments, this finding is analogous to, but not as extreme as, the variation in ATAC-seq reads obtained in different cells in single-cell experiments. For example, in the window shown in the middle panel of Figure 1B, the most highly methylated fiber has more than 10x as many methylated adenosines than the least-methylated fiber. If accessible adenosines were methylated at random with respect to fiber, we would expect a much tighter distribution of methyl-adenosines per fiber. This discrepancy can be quantified using a Q-Q plot in which the number of m6As observed per fiber is compared to the number of m6As expected per fiber, given that methylations occur at random with respect to fiber (**Fig 1E**, black dots). If we convert the number of m6As to z-scores for windows in different genomic locations, we can normalize for regional differences in accessibility and overlay multiple windows (**Fig 1E**, purple hexplot). The observed range of z-scores is at least 20x greater than that expected if m6As were distributed randomly across fibers. It remains unclear to us why the level of methylation is so variable across fibers at a single genomic location within a single sample, but we suspect that it is related to the fact that, to preserve the native chromatin state, nuclei must be handled very gently (i.e. cannot be mixed vigorously), and therefore the Hia5 enzyme may not be evenly distributed amongst the nuclei suspension.

Perhaps related to this per-fiber unevenness in methylation levels, we have found it difficult to obtain consistent methylation rates between different preparations of plant samples, even when using the same reaction conditions (e.g. units of methyltransferase, number of protoplasts, incubation time). This might be in part due to the extra care required to obtain nondegraded, high molecular weight DNA from plant tissue. Direct nuclei preps from frozen tissue have resulted in highly degraded DNA and from fresh tissue have limited scale due to the DNA loss during size selection and from additional clean up steps to remove excess polysaccharides and plant contaminants that can interfere with size selection and library preps. To obtain high quantities of quality plant DNA we protoplast, or enzymatically remove the wall from live cells, then chemically denature the plasma membrane and gently spin to release the nuclei (Bubb et al. 2025). Since we standardize the number of protoplasts prior to nuclei extraction, there may be more variation in the amount of DNA exposed to the methyltransferase between different Fiber-seq preps compared to the mammalian studies that directly extract and normalize the number of nuclei. We solve this problem by preparing multiple samples and evaluating sample quality on a pilot set of only tens of thousands of Fiber-seq reads before collecting the millions of reads necessary for a full analysis.

### 2. The effect of adenosine methylation rate and read depth on inferred fiber-level and genomic chromatin states

The Stergachis and Noble labs developed tools to infer multiple types of chromatin states from the raw adenosine methylation data. The fibertools pipeline (Jha et al. 2024) begins by using the positions of enzymatically methylated adenosines to infer nucleosome positions (unmethylated regions) and methylation sensitive patches (MSPs) on individual fibers. The FIRE pipeline (Vollger et al. n.d.) can then be used to label certain MSPs as Fiber-seq Inferred Regulatory Elements (FIREs) on individual fibers, with the remainder labeled as linkers between nucleosomes (**Fig 2A**).

**Figure 2.**
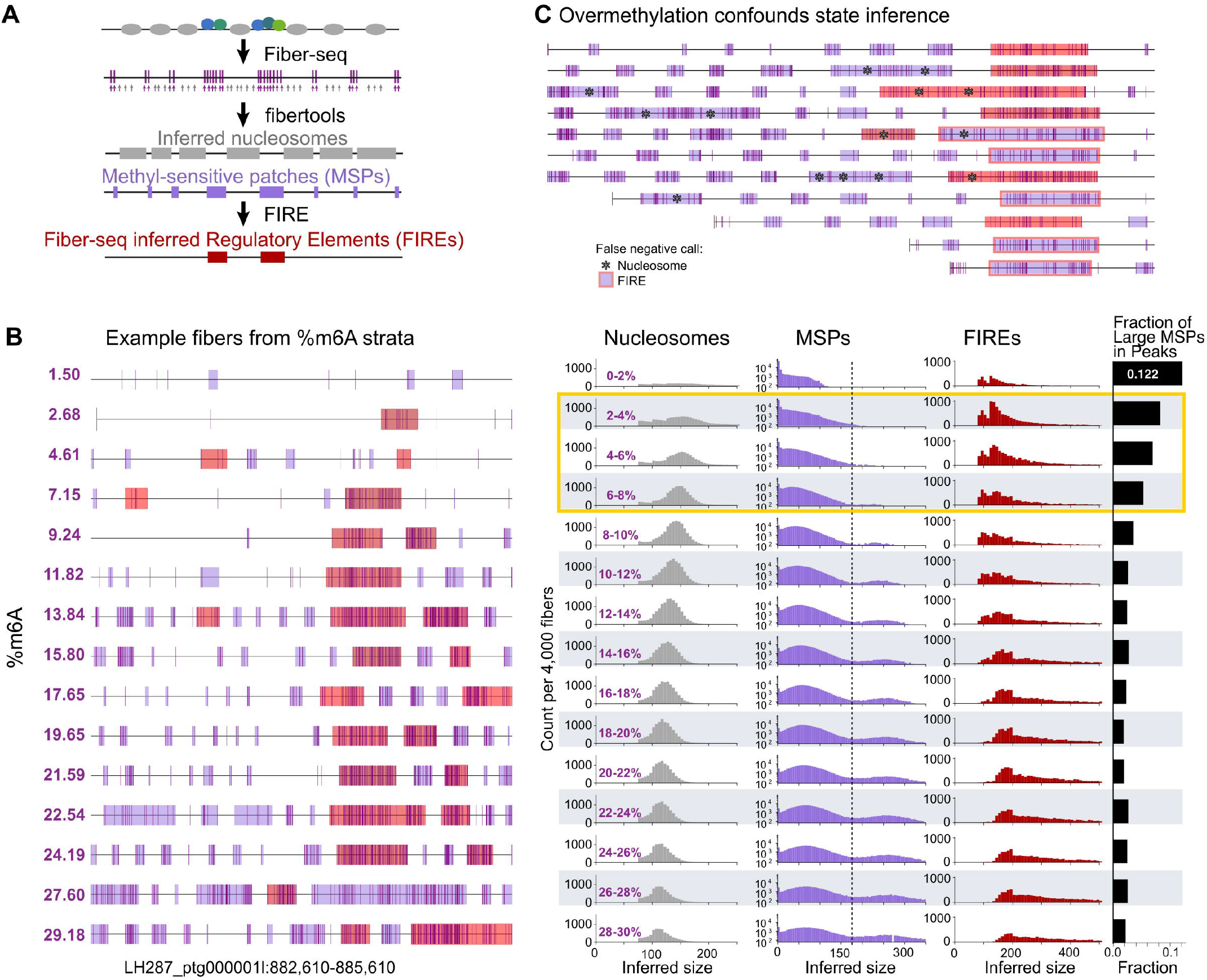
The effect of methylation levels on per-fiber state inference. **(A)** Illustration of the use of the fibertools and FIRE pipelines to infer the true chromatin state of a 4 kb section of a single fiber [LH244_ptg000011l:120,688,000-120,692,000]. True chromatin state is represented by the top-most graphic, with gray ovals indicating nucleosome-occupied nucleotides and blue and green circles indicating transcription-factor-occupied nucleotides. **(B)** *Example fibers from %m6A strata:* Read segments in a 3 kb region [LH287_ptg000001l:882,610-885,610] with varying %m6A. *Nucleosomes:* Histograms of nucleosome sizes, stratified by %m6A, range indicated in dark purple. The same number of fibers is in each bin, but the number of nucleosomes varies. *MSPs:* Histograms of MSP sizes, stratified by %m6A. The same number of fibers is in each bin, but the number of MSPs varies. *FIREs:* Histograms of FIRE sizes, stratified by %m6A. The same number of fibers is in each bin, but the number of FIREs varies. *Fraction of Large MSPs in Peaks:* Fraction of large (>=175 bp) MSPs that overlap high-quality FIRE peaks. **(C)** Example of overmethylation, resulting in long MSPs that contain methylated nucleosomes (false negative nucleosomes indicated with asterisks). These erroneously long MSPs confound FIRE-calling, resulting in underidentification of FIREs within FIRE peaks (false negative FIREs indicated by red outline around light purple MSP). [Og_chr01:891,000-893,000].

The genomic position of FIRE peaks (ACRs) is inferred from the combination of information provided by fibers underlying that location. Therefore, it is important to understand how methylation levels affect these inferred fiber-level states. For illustration, we display single fibers with varying methylation rates spanning a 3-kb region (**Fig 2B**, Example fibers from %m6A strata). We then plot the length distributions of inferred nucleosomes (**Fig 2B**, Nucleosomes), MSPs (**Fig 2B**, MSPs), and FIREs (**Fig 2B**, FIREs) on same-sized batches of DNA molecules (fibers) with differing percentages of adenosines methylated taken from a single sample preparation. As the percent of adenosines methylated increases, we observe (1) the inferred nucleosome sizes decrease, while MSP and FIRE sizes increase, (2) the emergence of a second mode of extra-large (>175-bp) MSPs, and (3) disappearance of the first mode of small FIREs (<125 bp in length), which have been shown to comprise a category of accessible regions largely undetected by ATAC-seq in a tissue-matched sample, yet coincide with ATAC-seq peaks in other tissues (Bubb et al. 2025).

As suggested in **Figure 2A**, some of the large (>175-bp) MSPs are converted to FIREs. However, especially in highly methylated fibers, many of the large MSPs are not reflective of genomic regions of accessibility but instead are cases where adenosine methylation occurs on DNA that is wrapped around a nucleosome, resulting in merging of linkers across nucleosomes (**Fig 2C**, long MSPs containing asterisks). These erroneously large MSPs become statistically conflated with large MSPs at true sites of accessibility and result in undercalling of true FIREs (**Fig 2C**, red outlined MSPs). To quantify the fraction of large MSPs in regions of true genomic chromatin accessibility (i.e. those that should be converted to FIREs), we calculated the Fraction of Large-MSP-bps in Peaks (FLiP score), using only well-supported FIRE peaks, having 50% or more underlying fiber-bps called as FIRE (**Fig 2B**, Fraction of Large MSPs in Peaks). To ensure high FLiP scores and to adequately detect biologically significant small FIREs, we conclude that fibers with 2-8% of adenosines methylated provide the most accurate and rich FIRE information.

### 3. The effect of adenosine methylation levels on peak-calling quantity and quality Next, we examine the factors affecting genomic-level FIRE peak-calling

To determine whether overall sample methylation rates affect the number of FIRE peaks called, we merged data from two samples that were treated with either low or high (**Fig 3A**, top and bottom, respectively) levels of methyltransferase. Using the same number of reads from each pooled set (4.27 M), the FIRE pipeline identified 66,889 FIRE peaks in the samples with low levels of methylation fewer than 3% more FIRE peaks (68,709) in the samples with high levels of methylation (**Fig 3B**, left), illustrating that the peak-calling feature of the FIRE pipeline is fairly robust to methylation distribution differences in samples. Furthermore, the quality of the peaks, defined by the distribution of FIRE accessibility scores per FIRE peak, were remarkably similar (**Fig 3B**, right)

Read depth, on the other hand, strongly affects both the number and quality of FIRE peaks called. This is expected for any significance-based peak caller, such as the commonly-used MACS2 for ATAC-seq (Gaspar 2018; Bubb & Deal 2020), however it is striking when quantified. We used downsampling to examine the relationship between read depth and number and quality (**Fig 3C**, top and bottom, respectively) of called FIRE peaks, with quality defined as the fraction of total underlying fiber bps labeled as FIREs (FIRE accessibility score). As expected, while the number of FIRE peaks increases with read depth, the fraction of FIRE peaks with low FA score increases, suggesting that above a certain depth, only transiently or partially accessible ACRs are identified -- along with a certain number of false positives. The reason for this, in short, is that FIRE peaks are called as a function of both the *quality*, reflected in degree of ‘redness’, and *number* of FIREs underlying that location and are, therefore, sensitive to local read depth.

**Figure 3.**
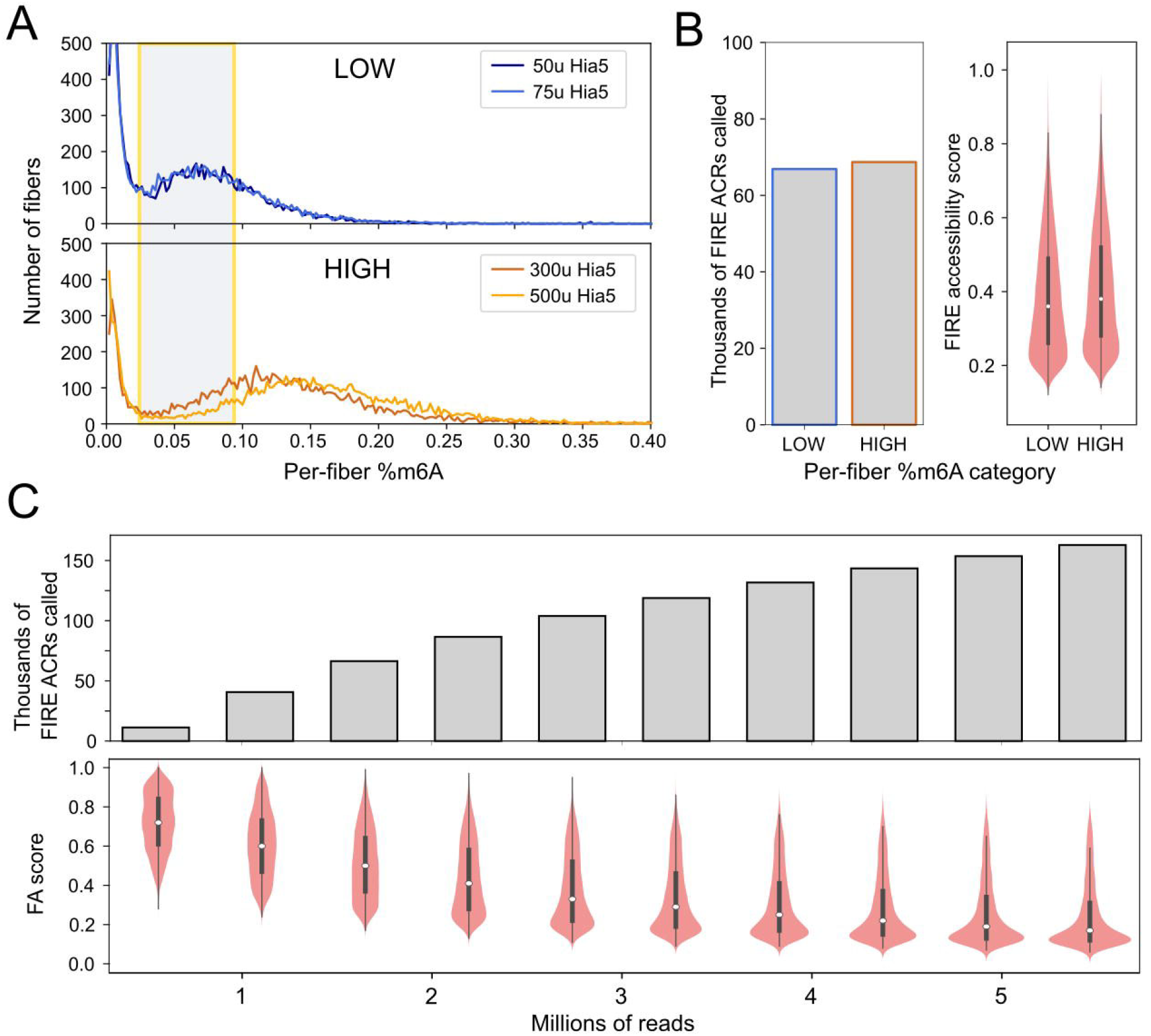
Effect of methylation levels and read depth on FIRE peak calling. **(A)** The distribution of percent adenosines methylated per fiber for two human cell line (GM12878) samples treated with a (top) low or (bottom) high amount of Hia5 enzyme units. The yellow box indicates the %m6A range recommended for per-fiber state feature inference. **(B)** The number of FIRE peaks called for low- and high-methylation samples (left). The FIRE accessibility score (fraction of fiber coverage called as FIREs) for each set of FIRE peaks (right). **(C)** The increase in the number of called FIRE peaks as read depth increases (top), and the mean FIRE accessibility score for called FIRE peaks decreases as read depth increases (bottom).

### 4. Comparing Fiber-seq results across divergent genomes

The above analyses establish a baseline of how the FIRE pipeline acts on samples of varying methylation levels and read depths. We next turn to how best to compare across samples with different genomes.

Although Fiber-seq was initially developed in Drosophila (Stergachis et al. 2020), much of the more recent use has been in human tissue. Our lab has further expanded the use of Fiber-seq to maize (Bubb et al. 2025), and we are now expanding to other plant species as well as multiple within-species strains and their hybrids. With this data we can compare accessibility across homologous regions as well as examine the effects of trans-environments on accessibility for identical underlying sequences, specifically how genome accessibility changes in an F1 context. Maize and humans have very different evolutionary histories, but there are certain similarities. Humans have experienced repeated bottleneck and founder effects (Tournebize et al. 2022).

Maize, as a domesticated species, has also undergone several population bottlenecks, and yet domesticated maize retains a surprising level of genetic diversity. This diversity is at least partially due to active transposition, leading to transposon-level presence/absence polymorphisms and larger-scale structural variation (Buckler et al. 2006; Munasinghe et al. 2023), as well as persistent introgression by its wild ancestor, teosinte (Yang et al. 2023). The suggestions we propose below were developed for maize but are generally applicable for any cross-genome comparisons.

#### First, when collecting Fiber-seq reads from hybrid individuals, we can only compare accessibility of the haplotypes in regions where the Fiber-seq reads are uniquely mappable

To ensure that fibers are mapped to their correct genome of origin, we align our reads to a compound genome consisting of both parental genomes concatenated and maintain only uniquely mappable reads (mapq=60 using pbmm2). For consistency, we also map fibers from parental lines to this compound genome, even though we know that all fibers originate from this genome. We determined the fraction of the genome that is uniquely mappable by our long-reads by simulating reads from an *in silico* F1 genome, using read lengths and per-base error distributions empirically derived from previous Fiber-seq reads, and mapping them to either parental-only or compound genomes. We then report the percent of the genome uniquely mappable for each parent genome with either mapping strategy (**Fig 4A**). For all maize strain pairs examined, only 70-80% of either parental genome is uniquely mappable by F1 reads to an F1 (compound) genome. Note that this fraction can vary for the same parent when it is paired with a different parent, depending upon how similar the genomes of the two co-parents are. For example, LH244 has better alignment with its parental genome when concatenated to Ki3 vs. Oh43. In contrast, more than 95% of the genome is mappable if simulated reads are mapped to parental-only genomes rather than compound genomes.

**Figure 4.**
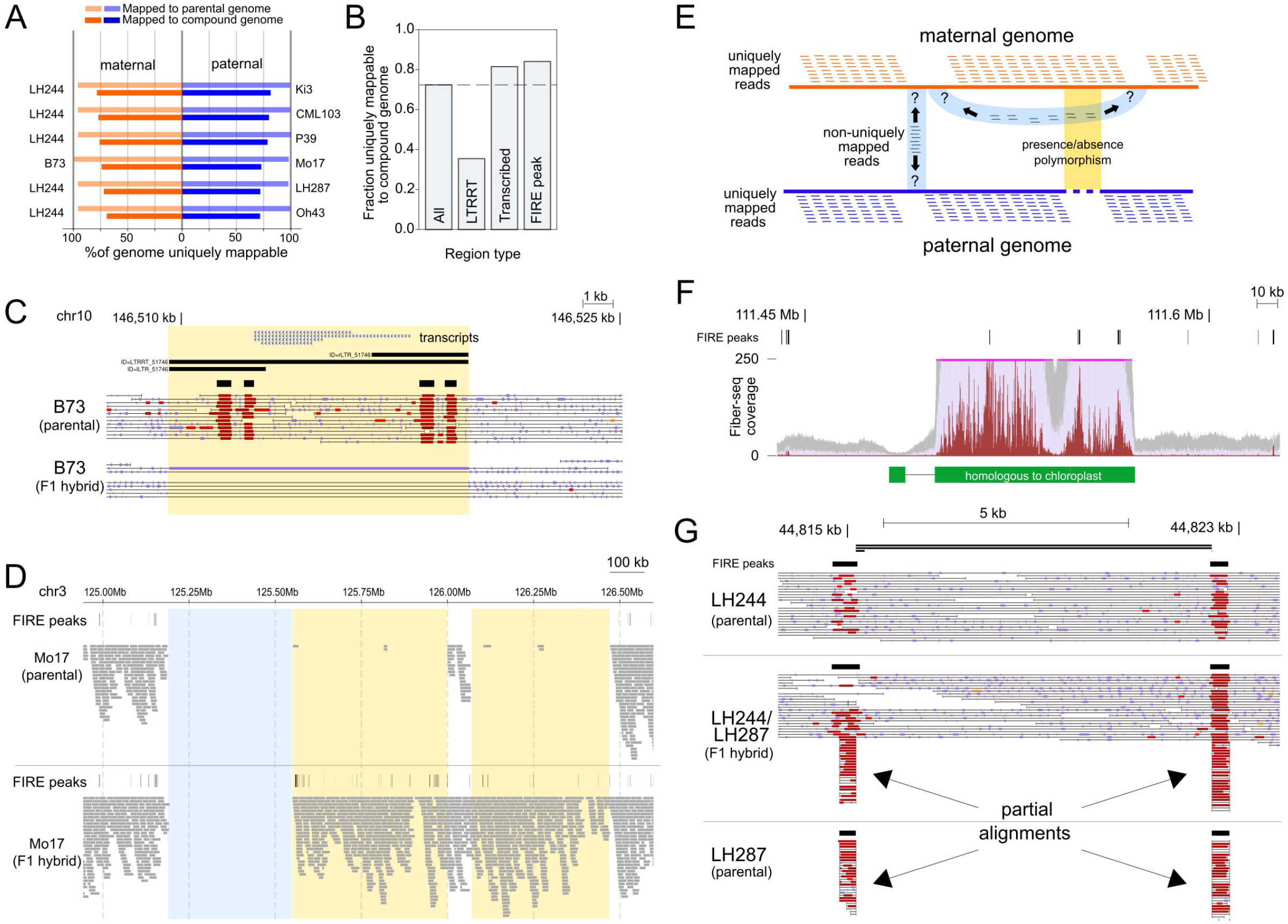
Mapping errors that can result in miscalling of differentially accessible regions. **(A)** Each row reports the fraction of maternal and paternal genomes that are uniquely mappable by Fiber-seq-like simulated reads, when aligning to either a parent-only genome (light color) or a compound genome (dark color). **(B)** Bar plot of fraction bps in regions uniquely mappable to a compound genome: (i) all bps in genome (ii) bps annotated as LTR retrotransposons, (iii) bps annotated as transcribed, and (iv) bps annotated as FIRE peaks when fibers are mapped to a single genome. **(C)** A transposon present in parental B73 and not the B73 genome in our B73xMo17 F1 is highlighted in yellow (chr10:146,505,085-146,527,310). **(D)** Large region with coverage in parental Mo17 but not the F1 Mo17 haplotype (chr3:124,837,001-126,845,980). Yellow highlighted regions indicate sequence missing from the Mo17 genome that is in our sequenced F1. Blue highlighted regions indicate regions where reads cannot be uniquely mapped. **(E)** Cartoon showing distinction between non-uniquely-mappable regions (blue) and regions missing relative to mapped-to genome (yellow). **(F)** An example of a region within the nuclear genome that has homology to the chloroplast genome. Fiber-seq reads originating from the chloroplast genome are misaligned to the nuclear location [LH287_ptg000001l:111,375,000-111,625,000]. **(G)** An example of FIRE peak calling artificially enhanced by Fiber-seq reads from one haplotype by partially aligning to the wrong haplotype in the F1 (LH244_ptg000012l:44,811,956-44,824,088).

We then determined whether different types of base pairs were more likely to be in uniquely mappable regions, looking specifically at LH287 in the LH244 x LH287 parental pair. While 76% of all genomic bps were uniquely mappable, less than 40% of LTR retrotransposon bps were in uniquely mappable regions. Transcribed or accessible (FIRE peak) bps were more likely to be in uniquely mappable regions as compared to all genomic bps (**Fig 4B**).

#### Second, presence/absence polymorphisms, such as transposon insertions, structural variation, or copy number variation, should not be mistaken for accessibility differences

We soon noticed that although the genomes in our sequenced inbred lines and F1s were nominally the same (e.g. P1=B73; F1=B73xMo17; P2=Mo17) there were genomic presence/absence differences between them. For example, an LTR retrotransposon is present in the mapped-to reference B73 (v5) and the sequenced parental B73 but not in the F1 B73 genome (**Fig 4C**, yellow highlighted region). In some cases, regions missing in the parental version, but not the F1 version, of the Mo17 genome were quite large -- on the scale of megabases (**Fig 4D**, yellow highlighted region). Note that regions lacking coverage in both the parental and F1 Mo17 genomes are likely not uniquely mappable in the B73xMo17 compound genome (**Fig 4D**, blue highlighted region). In general, if the specific strain that we sequence is missing genetic material present in the mapped-to reference sequence, there will be no Fiber-seq coverage of those regions. Thus, there are two broad reasons for missing coverage: (i) lack of unique mappability, and (ii) the sequenced genome is missing a part of the mapped-to genome (**Fig 4E**, blue and yellow shading, respectively). Such regions should not be included in lists of differential accessibility.

#### Third, reads should be aligned to genomes that include organellar genomes, to prevent ectopic alignment

Substantial stretches of the nuclear genome have sequence homology to organellar genomes, and fibers originating from organellar genomes should be allowed to align to their true genome of origin. Organellar genomes typically have completely accessible chromatin and are present at much higher copy number than the nuclear genome. Ectopic mappings can introduce both (i) false positives, because the accessibility and coverage of fibers matching such regions are much higher than nuclear-derived fibers, and (ii) false negatives, because this region may indeed contain accessible regions that should not be ignored or discarded. For example, when we align Fiber-seq reads to only the nuclear genome, regions with homology to chloroplast genome have more than 10x higher coverage than the rest of the genome and are correctly flagged as regions of “unreliable coverage” by the FIRE pipeline (**Fig 4F**). To prevent this erroneous mapping, the chloroplast and mitochondrial genomes should be appended to the compound nuclear genomes before alignment, allowing fibers originating from the organellar genomes to uniquely map to their correct location.

#### Fourth, partial alignments should be removed prior to running the FIRE pipeline, as they can introduce another source of artifact to both FIRE peak calling and FIRE accessibility score quantification

For example, sections of fibers deriving from the parental LH287 genome align to regions immediately flanking an intact LTR retrotransposon in the parental LH244 genome (**Fig 4G**). Such alignments can and should be removed early in the analysis pipeline.

### 5. Detecting differentially accessible FIRE peaks

Finally, after appropriately normalizing and controlling for artifacts, we can begin looking for truly differentially accessible FIRE peaks across samples.

FIRE peak calling is a useful way to identify the small fraction of the genome that is above some threshold of accessibility. However, among regions identified as FIRE peaks, there are different degrees and types of accessibility. For example, some called FIRE peaks cover more contiguous bps than others. Some FIRE peaks consist of smaller per-fiber FIREs that don’t occupy exactly the same sequence, which can result in large FIRE peaks. Some FIRE peaks have a smaller fraction of underlying fibers that contain FIREs. We are only beginning to understand the significance of these differences. Likewise, when identifying differential FIRE peaks, we should use metrics that are sensitive to these state differences and not simply rely on peak presence/absence.

To illustrate these points, we are comparing accessible regions between the B73 genome in the parental inbred context and in an F1 (B73 x Mo17) context. However, a similar approach could be used to compare accessibility between homologous regions in B73 and Mo17 parentals or across any pair of genomes. The difference between genomes at each union FIRE peak can be described with an effect size and a significance score. Using both the effect size (absolute difference in either per-base or per-fiber FIRE proportions between compared samples) and significance (p-value from a two-proportion Z-test, comparing either the proportion of bps called as FIREs or the proportion of fibers containing FIREs in one sample vs the other), we can identify a spectrum of differential accessibility across samples at each peak (**Fig 5A,B**). Effect size and significance, while correlated, do not provide identical information. Three examples of FIRE peaks, each shown within their 3-kb window contexts, are displayed to highlight the features that can affect these four metrics.

**Figure 5.**
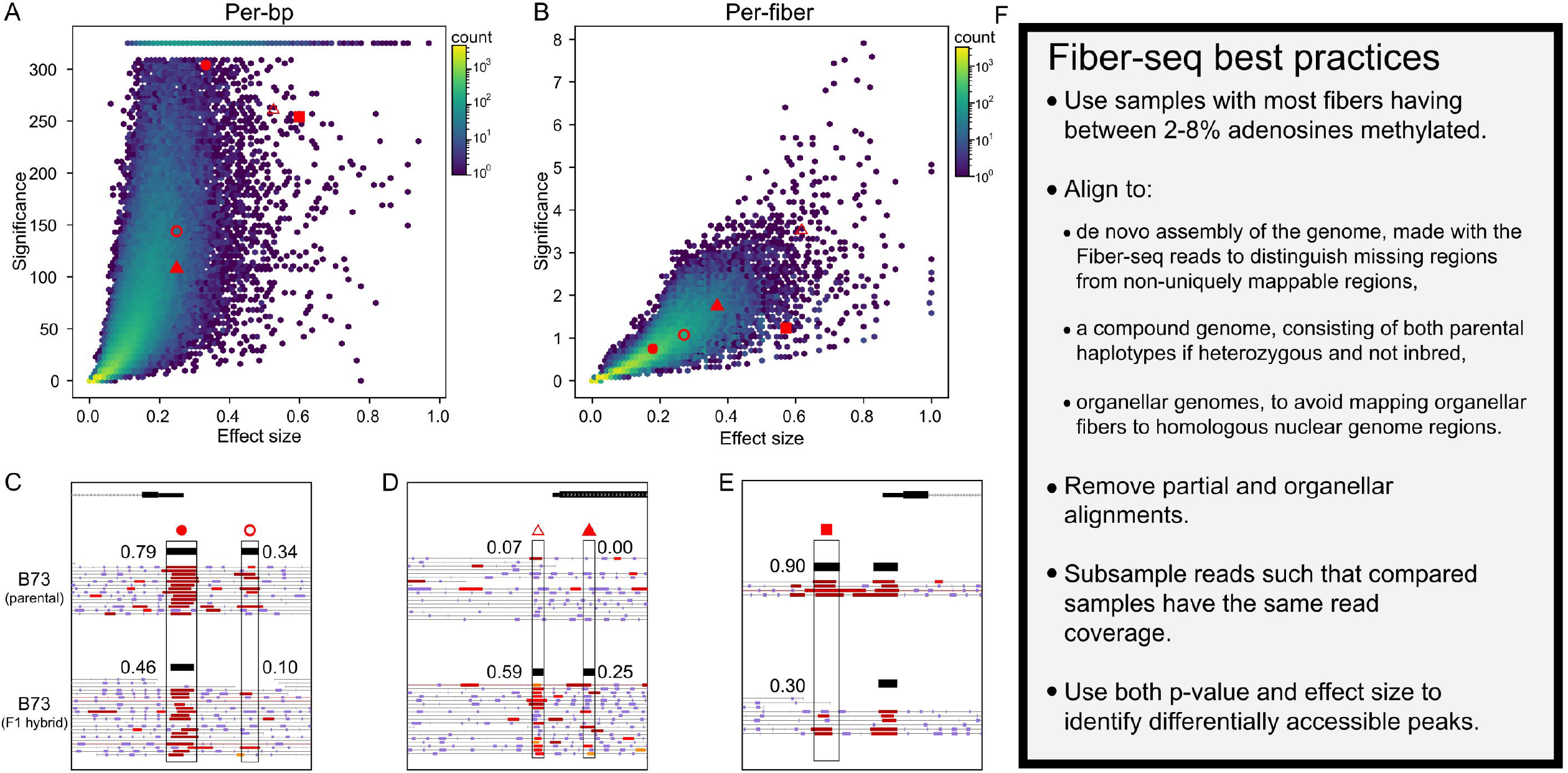
Identifying true positive differentially accessible regions. **(A)** X-axis: Effect size (absolute between-sample difference in FIRE accessibility score); y-axis: Significance [-log10(p-value)] based on a comparison of the proportions of bps tagged as FIRE element. The total number of peaks (<=500bp) is 77,297. Red dots are union peaks shown in C-E. **(B)** X-axis: Effect size (absolute between-sample difference in number of fibers containing a FIRE); y-axis: Significance [-log10(p-value)] based on a comparison of the proportion of fibers containing a FIRE. The total number of peaks (<=500bp) is 77,294. Red dots are union peaks shown in C-E. **(C-E)** Numbers next to union peak (rectangle) indicate FIRE Accessibility score for that sample. **(C)** A moderate loss of accessibility in the F1 context relative to the inbred context (solid red circle). Presence/absence peak (open red circle). Window: [B73_chr03:227,051,500-227,054,500; gene: Zm00001eb160680]. **(D)** Gain of accessibility in the F1 context relative to the inbred context (solid red triangle). Presence/absence peak (open red triangle). Window: [B73_chr01:36,742,000-36,745,000; gene: Zm00001eb011320]. **(E)** Loss of accessibility in the F1 context relative to the inbred context but low read coverage (solid red square). [Window: B73_chr03:187,818,500-187,821,500; gene: Zm00001eb148930]. **(F)** A summary of analytic guidelines.

First, union peaks with high fiber-bp coverage (many fibers, long union peak) and differences primarily in the size of called FIREs will have moderate per-bp and per-fiber effect sizes and high significance in the per-bp proportions test but not in the per-fiber proportions test (**Fig 5A,B,C**). Additionally, such regions are likely to be identified as FIRE peaks in both samples. Intermediate accessibility levels may reveal a snapshot of the transition from accessible to inaccessible, or vice versa. For example, we previously used Fiber-seq to identify FIRE peaks within long terminal repeat retrotransposons (LTRRTs). LTRRTs with two peaks in each LTR, putative enhancers and promoters, had higher per-bp accessibility and low rates of CpG methylation, whereas LTRRTs with only a single peak per LTR had both lower per-bp accessibility and higher rates of CpG methylation (Bubb et al. 2025).

Second, small union peaks with a high proportion of fibers containing FIRE in one sample and not the other will attain a high significance score in the per-fiber proportions test. The per-fiber test statistic is not affected by the size of the peak, so excels at finding differences between narrow peaks (**Fig 5A,B,D**). Third, union peaks with low fiber coverage will have low significance by the per-fiber proportions test but may have higher significance by the per-bp proportions test if the peak is large (**Fig 5A,B,E**).

Finally, and perhaps most importantly, peak presence/absence is generally a poor method of identifying regions with true accessibility difference between samples, even for samples with equal read depth. For example, two such regions are present in our windows (**Fig 5C&D**, open circle and open triangle). They have low values for all four metrics (**Fig 5A&B**). Therefore, it is important to critically assess differential peak types to fully harness the power of Fiber-seq in identifying differential biological contexts rather than oversimplifying to binary peak calling.

In summary (**Fig 5F**), we recommend comparing samples with similar per-fiber rates of adenosine methylation, ideally with the majority of fibers having a percent of adenosines methylated within the range of 2-8%. In our hands, it has been useful to prepare multiple samples and sequence them at low coverage in order to determine if the adenosine methylation rates and nucleosome size distributions are within the optimal range before obtaining the full coverage. Next, if analyzing hybrids, align to a compound genome that consists of both parental genomes, ideally assembled *de novo* from the Fiber-seq data itself, rather than a published genome, as well as any plastid genomes (i.e. chloroplast and mitochondrial genomes. Remove partial and organellar alignments, then randomly subsample reads such that each sample has the same genomic coverage, with the hybrid sample having roughly twice as many reads as either parent to obtain similar coverage over each parental genome. Next run fibertools (Jha et al. 2024) to infer nucleosomes and MSPs, and then the FIRE pipeline (Vollger et al. n.d.) to call FIREs on individual fibers and FIRE peaks at genomic locations. Generate a set of union FIRE peaks by merging peaks called on each sample to compare (e.g. using bedops (Neph et al. 2012) or bedtools (Quinlan 2014)). Finally, define thresholds for effect size (e.g. absolute difference in FIRE accessibility) and p-value (e.g. using a two-proportion Z-test) that identify the type of differential accessibility that is of interest to you.

## Acknowledgements

We thank Bing Yang (University of Missouri) for rice samples, Cinta Romay (Cornell University) for maize seeds, Mitchell Vollger (University of Utah), Shane Neph and the Stergachis Lab (University of Washington), Bryan Ramirez-Corona and the Queitsch/Cuperus Lab (University of Washington) for helpful discussions, and Andrew Stergachis (University of Washington) for the human cell line data.

This work was supported by the National Science Foundation (IOS Division of Integrative Organismal Systems grant no. 2519784 to C.Q. and PlantSynBio grant no. 2240888 to C.Q.), and the National Institutes of Health (T32 training grant no. HG000035 to J.T., NIGMS grant no. R01-GM079712 to J.T.C. and C.Q., and NIGMS MIRA grant no. 1R35GM139532 to C.Q.).

## References

Baek, S. & Lee, I., 2020. Single-cell ATAC sequencing analysis: From data preprocessing to hypothesis generation. Computational and Structural Biotechnology Journal, 18, pp.1429–1439.

Bubb, K.L. et al., 2025. The regulatory potential of transposable elements in maize. Nature plants, 11(6), pp.1181–1192.

Bubb, K.L. & Deal, R.B., 2020. Considerations in the analysis of plant chromatin accessibility data. Current opinion in plant biology, 54, pp.69–78.

Buckler, E.S., Gaut, B.S. & McMullen, M.D., 2006. Molecular and functional diversity of maize. Current Opinion in Plant Biology, 9(2), pp.172–176.

Deal, R.B. & Henikoff, S., 2011. The INTACT method for cell type-specific gene expression and chromatin profiling in Arabidopsis thaliana. Nature protocols, 6(1), pp.56–68.

Gaspar, J.M., 2018. Improved peak-calling with MACS2. bioRxiv, p.496521. Available at: https://www.biorxiv.org/content/10.1101/496521v1 [Accessed August 7, 2019].

Groza, C. et al., 2023. Genome graphs detect human polymorphisms in active epigenomic state during influenza infection. Cell Genomics, 3(5), p.100294.

Hu, Y. et al., 2025. Multiscale footprints reveal the organization of cis-regulatory elements. Nature, 638(8051), pp.779–786.

Jha, A. et al., 2024. DNA-m6A calling and integrated long-read epigenetic and genetic analysis with fibertools. Genome Research, 34(11), pp.1976–1986.

Marand, A.P. et al., 2025. The genetic architecture of cell type-specific cis regulation in maize. Science (New York, N.Y.), 388(6744), p.eads6601.

Munasinghe, M. et al., 2023. Combined analysis of transposable elements and structural variation in maize genomes reveals genome contraction outpaces expansion. PLoS genetics, 19(12), p.e1011086.

Neph, S. et al., 2012. BEDOPS: high-performance genomic feature operations. Bioinformatics, 28(14), pp.1919–1920.

Quinlan, A.R., 2014. BEDTools: The Swiss-army tool for genome feature analysis: BEDTools: The Swiss-army tool for genome feature analysis. Current Protocols in Bioinformatics, 47(1), pp.11.12.1–34.

Stergachis, A.B. et al., 2020. Single-molecule regulatory architectures captured by chromatin fiber sequencing. Science, 368(6498), pp.1449–1454.

Swift, J. et al., 2023. Single nuclei sequencing reveals C 4 photosynthesis is based on rewiring of ancestral cell identity networks. Plant Biology. Available at: https://www.biorxiv.org/content/10.1101/2023.10.26.562893v1.full.pdf.

Tournebize, R., Chu, G. & Moorjani, P., 2022. Reconstructing the history of founder events using genome-wide patterns of allele sharing across individuals. PLoS Genetics, 18(6), p.e1010243.

Vollger, M.R. et al., A haplotype-resolved view of human gene regulation. Available at: 10.1101/2024.06.14.599122.

Vollger, M.R. et al., 2025. Synchronized long-read genome, methylome, epigenome and transcriptome profiling resolve a Mendelian condition. Nature Genetics, 57(2), pp.469–479.

Wang, C. et al., 2025. Computational analyses and challenges of single-cell ATAC-seq. Genomics, Proteomics & Bioinformatics, 23(6), p.qzaf115.

Yang, N. et al., 2023. Two teosintes made modern maize. Science, 382(6674), tp.eadg8940.

